# Pinwheel-dipole configuration in cat early visual cortex

**DOI:** 10.1101/009308

**Authors:** Jérôme Ribot, Alberto Romagnoni, Chantal Milleret, Daniel Bennequin, Jonathan Touboul

## Abstract

In the early visual cortex, information is processed within functional maps whose layouts are thought to underlie visual perception. However, the precise organization of these functional maps as well as their interrelationships remains largely unknown. Here, we show that spatial frequency representation in cat early visual cortex exhibits singularities around which the map organizes like an electric dipole potential. These singularities are precisely co-located with singularities of the orientation map: the pinwheel centers. To show this, we used high resolution intrinsic optical imaging in cat areas 17 and 18. First, we show that a majority of pinwheel centers exhibit in their neighborhood both semi-global maximum and minimum in the spatial frequency map, contradicting pioneering studies suggesting that pinwheel centers are placed at the locus of a single spatial frequency extremum. Based on an analogy with electromagnetism, we proposed a mathematical model for a dipolar structure, accurately fitting optical imaging data. We conclude that a majority of orientation pinwheel centers form spatial frequency dipoles in cat early visual cortex. Given the functional specificities of neurons at singularities in the visual cortex, it is argued that the dipolar organization of spatial frequency around pinwheel centers could be fundamental for visual processing.

**ABBREVIATIONS:**

OR: Orientation
SF: Spatial frequency
PC: Pinwheel center
A17: Area 17
A18: Area 18

## 1 INTRODUCTION

The brain contains the representations of multiple sensory features that form continuous and orderly maps. These maps subtend perception, yet the principles governing their organization remain fundamentally unknown. Uncovering the fine structure of maps can reveal their underlying organizing principles and shed new light on how the brain processes the sensory information.

The early visual cortex (V1 and V2) of higher mammals provides a particularly appropriate cortical area to investigate these questions. Its neurons are selective locally to several attributes of the visual scene, such as the orientation (OR) of a stimulus (Hubel & Wiesel, 1959) and its spatial frequency (SF) (Movshon et al., 1978). Neurons with similar functional properties are clustered vertically and form functional maps parallel to the cortical surface. The organization of the OR map has been extensively studied in the past 20 years: it consists of regular zones where preferred OR varies smoothly, together with singularities, the *pinwheel centers* (PCs), in the vicinity of which all ORs are represented (Bonhoeffer & Grinvald, 1991).

In contrast, the existence and the organization of the SF map have been the subject of long-standing debates over the last decades. Evidence however accumulates that SF is organized in a continuous manner in several mammals, including cat (Issa et al., 2000; Villeneuve et al., 2009; Tani et al., 2012; Ribot et al., 2013) and monkey (Nauhaus et al., 2013). Yet, how the layout of the SF map is constrained to the OR map and how these constrained might subtend visual perception remain disputed. In cat, for instance, pioneering investigations have suggested that OR PCs are situated preferentially at local SF extrema (Shoham et al., 1997; Issa et al, 2000; Issa et al., 2008). These results fit with the uniform coverage hypothesis (Swindale et al, 2000) according to which all combinations of the attributes should be uniformly represented in the cortex. On the other hand, it was recently argued that these findings might be the result of high-pass filtering methods leading to extreme SF values being over-representation at PC (Ribot et al., 2013). Instead, these new data based on recent intrinsic optical imaging techniques (Kalatsky & Stryker, 2003) have rather suggested that SF gradients were sharp in these cortical locations (Ribot et al., 2013), arguing against uniform coverage hypothesis at PCs.

In this study, we reexamined this issue, and used high-resolution intrinsic optical imaging in cat visual cortex area 17 (A17) and area 18 (A18) to map the SF structure near OR PCs. First, we show that at a majority of pinwheel locations, the SF representation covers a wide range of SFs, with values including semi-global minima and maxima found over a hypercolumn. This intriguing property prompted a more accurate description of the structure of the SF representation. To this purpose, we introduced a mathematical model for a dipolar structure and show that it could be accurately fitted to the SF map near a majority of PCs. The functional role of this pinwheel-dipole configuration is then discussed and it is proposed that representing a large range of SFs close to PCs is beneficial for visual perception.

## 2 MATERIALS AND METHODS

Five adult cats (two males, three females), aged between 24 and 72 weeks and born from different litters in our colony, were studied. They were all in good health and had no apparent malformations or pathologies. All experiments were performed in accordance with the relevant institutional and national guidelines and regulations i.e. those of the Collège de France, the CNRS and the DDPP (JO 87-848, consolidated after revision on May 30, 2001, Certificate n° 75-1754, French “Ministère de l’Agriculture et de la Pêche”). They also conformed to the relevant regulatory standards recommended by the European Community Directive (Directive 2010/63/UE) and the US National Institutes of Health Guidelines.

### 2.1 Surgical procedure

On the day of the optical imaging experiment, animals were anaesthetized with Saffan® (initial, i.m., 1.2 mg/Kg; supplements, 1:1 in saline, i.v. as needed). After tracheal and venous cannulation, electrocardiogram, temperature and expired CO_2_ probes were placed for continuous monitoring. Animals were installed in a Horsley-Clarke stereotaxic frame and prepared for acute recordings. The scalp was incised in the sagittal plane, and a large craniotomy was performed overlying areas 17 and 18 of both hemispheres. The nictitating membranes were then retracted with eye drops (Neosynephrine® 5%, Ciba Vision Ophthalmics, France) and the pupils dilated with atropine eye drops (Atropine 1%, MSD-Chibret, France). Scleral lenses were placed to protect the cornea and focus the eyes on a tangent screen at a distance of 28.5 cm. The size of the lenses was adapted to the eye of each cat. Animals were then paralyzed with an infusion of Pavulon (0.2 ml/kg, i.e., 0.4 mg/kg i.v.) and breathing was assisted artificially through a tracheal cannula with a 3:2 mixture of N_2_O and O_2_ containing 0.5–1.0% isoflurane. The respiration frequency was adjusted to around 18 per minute and the volume adapted to the ratio of exhaled CO_2_ (pCO_2_ was maintained at 4%). Paralysis was maintained throughout the recording by continuous infusion of a mixture of Pavulon (0.1 ml/kg/h) diluted in glucose (5%) and NaCl (0.9 g/l). At the end of the recording session, the animal was given a lethal dose of pentobarbital. The experiments lasted less than 12 hours.

### 2.2 Optical imaging

The cortex was illuminated at 545 nm to reveal the vascular pattern of the cortical surface and at 700 nm to record the intrinsic signals. The focal plane was adjusted to 500 μm below the cortical surface. The optic discs were plotted by tapetal reflection on a sheet of paper covering the tangent screen. The centre of the screen was situated 8 cm (∼15°) below the middle of the two optic discs, that is, ∼8.5° into the lower visual field (Bishop et al., 1962). Intrinsic optical signals were recorded while the animal was exposed to visual stimuli displayed on a CRT monitor. The size of the display subtended a visual angle of ∼75° × 56°. The image had a resolution of 14 pixels per degree and the refresh rate was 88 Hz. Frames were acquired by CCD video camera (1M60, Dalsa, Colorado Springs, USA) at the rate of 40 frames per second and were stored after binning by 2 × 2 pixels spatially and by 12 frames temporally using the LongDaq system (Optical Imaging Inc., New York, USA). In three animals, images were acquired at a resolution of approximately 15.3 μm/pixel via a 50x50 mm tandem lens configuration. To illustrate the fine representation of SF around OR PC (Fig. 5 and Fig. S2), a 135×50 mm configuration was used to reach a spatial resolution of 5.9 μm/pixel.

### 2.3 Stimulation

Full-screen visual stimuli were presented continuously to the animal. Each stimulus consisted of sine-wave gratings drifting in one direction and rotating in a counter-clockwise manner (Kalatsky & Stryker, 2003). The angular speed of the rotation was 2 rotations per min. The temporal frequency of the drift was 2 Hz (Khaytin et al., 2008, Ribot et al., 2008). The contrast was set at 50% to ensure the production of smooth sine wave gratings (Xu et al., 2007). Thirty SFs ranging linearly in a logarithmic scale from 0.039 to 3.043 cycle/degree (cpd) were presented in random order. Ten full rotations were presented for each SF. At the end of the last rotation, the first frame of the first rotation for the next SF was presented without interruption. The total duration of the recording was 2.5 hours.

### 2.4 Image processing

Data analysis was based on the method developed by Kalatsky and Stryker (2003) to extract periodic signals from intrinsic data in Fourier space. First, data were pre-processed to remove slow-varying components. A common approach for this is high-pass filtering of images (Vanni et al., 2010). However, since SF representation is anisotropic in both A17 and A18, active domains may have very different sizes depending on the stimulus SF. High-pass filtering is thus risky since it spatially limits the size of the activated areas. Here, we used a multivariate analysis technique, the generalized indicator function method (Yokoo et al., 2001; Ribot et al., 2006), that does not require any assumptions about the spatial characteristics of the activity patterns.. This procedure was applied to raw data for each SF separately. A low-pass filter with a Gaussian kernel of around 30 μm half width was also applied for smoothing the data.

To construct the OR map, a Fourier transform was performed on the temporal signal of each pixel for all SFs together. The phase at the frequency of rotation was calculated to obtain the preferred OR (plus the hemodynamic delay) at each pixel (Kalatsky and Stryker, 2003). The hemodynamic delay (approximately 2.5 s) was subtracted from all OR maps.

Then intrinsic signals related to each SF were considered separately. For each pixel, the modulation of the signal induced by the rotation of the gratings was interpolated via a least-square method with a cosine function whose phase was equal to the preferred OR at this pixel and whose frequency was equal to half the frequency of rotation. As a result, magnitude maps for preferred ORs were obtained for each stimulus SF. These maps are referred as “maximal intensity maps”. Pixels with negative values, which corresponded to interpolation peaking at orthogonal ORs, were rectified to zero. Then, values were rescaled for each pixel by setting the greatest value among all SFs to 100 (Fig. S1).

Finally, the location of the functional border between A17 and A18 was estimated based on the difference between the maximal intensity maps at 0.15 cpd and 0.5 cpd (Bonhoeffer et al., 1995; Ohki et al., 2000).

### 2.5 Determination of the preferred spatial frequency

To evaluate the preferred SF at each pixel, the intrinsic signals were interpolated with a difference of Gaussians (DOG) function:

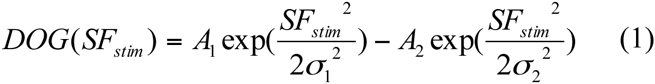

where *SF*_*stim*_ represents the SF of stimulation, *A*_1_ and *A*_2_ are the amplitude of each Gaussian function, and *σ*_1_ and *σ*_2_ are related to the bandwidth of each Gaussian function. Values for *A*_1_, *A*_2_, *σ*_1_ and *σ*_2_ were optimized to provide a least square error fit to the data using Matlab®.

At each pixel, the error-of-fit was defined by

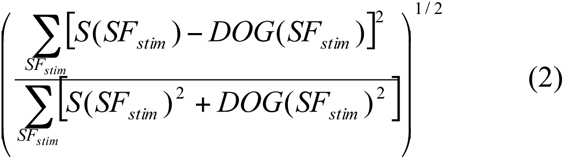

Where *S*(*SF*_*stim*_) represents the values of the intrinsic signal for the SF of stimulation. It ranged between 0 (perfect fit) and 1 (worst fit). Eventually, the preferred SF was extracted from the peak of the DOG tuning curve.

Examples preferred SF maps are shown in Fig. 1A for two different animals (White dots represent PC position). Pixels with an error-of-fit greater than 0.5 were discarded (black areas, Ribot et al., 2013). We took special care of the low-pass filtering and checked that it did not affect substantially the structure of these maps, particularly near PCs. As shown in Fig. 1B, the two maps remained highly stable even when low-pass filtering was discarded: the main structures and organization are conserved. Quantitatively, we showed (Fig. 1C) that it did not significantly change the preferred SF, neither near (<150 μm) nor far from PCs (>150 μm; linear regression, R^2^ > 0.98 in each case).

**Figure 1.**
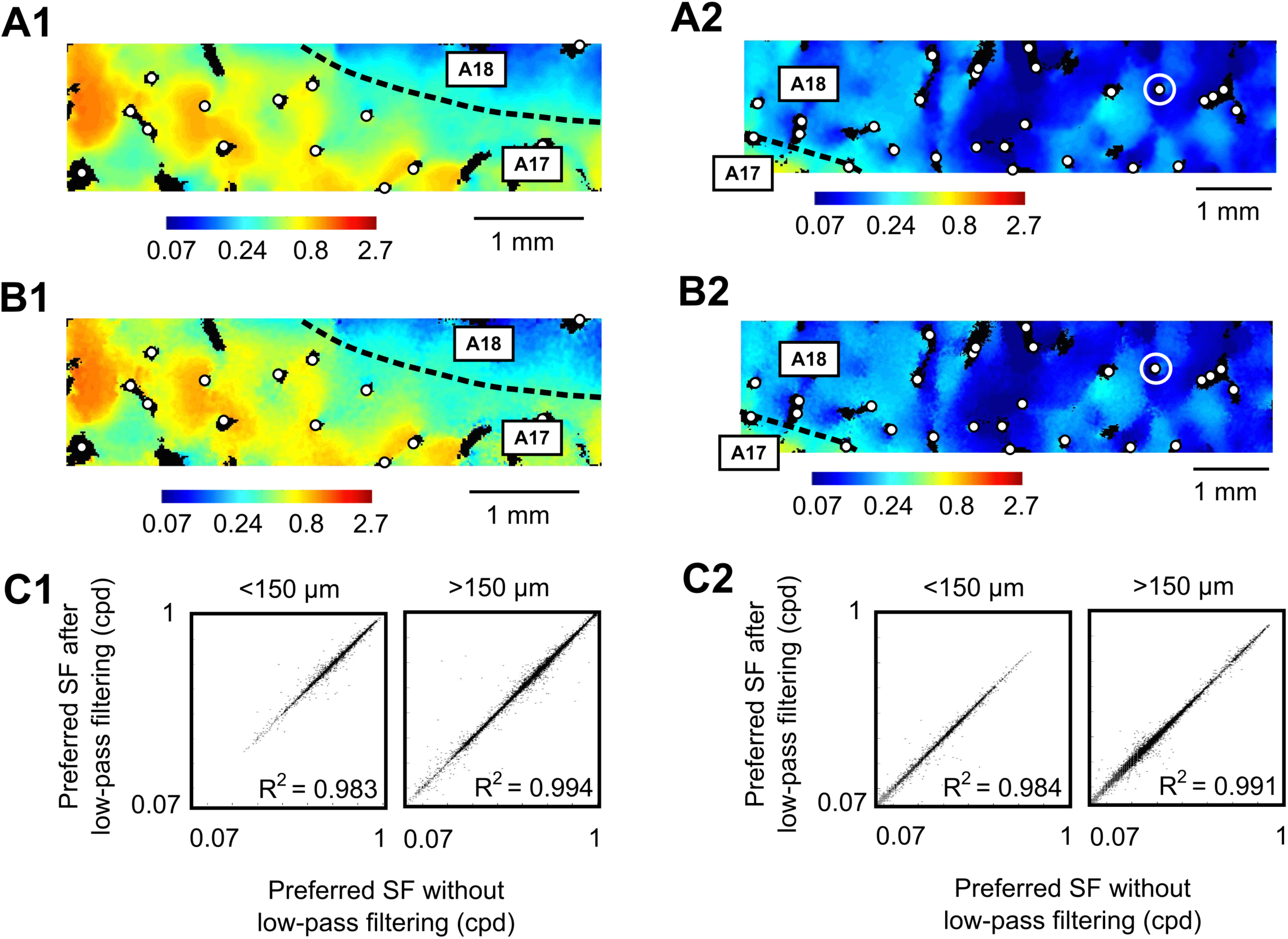
SF maps and low-pass filtering. (A) Two preferred SF maps corresponding to two different animals. White dots correspond to PC locations. Black areas have an error-of-fit > 0.5. The tuning curves of the PC circled in white in (A2) are represented in Fig. 2. (B) Same as (A) when low-pass filtering was discarded: the filtering did not induce any visible bias on the representation of preferred SF. (C) Correlation between maps in (A) and (B) for cortical areas near PCs (<150 μm) and far from PCs (>150 μm). Each dot corresponds to one pixel in the maps. The coefficient of determination R^2^ following linear regression was typically > 0.98.

**Figure 2.**
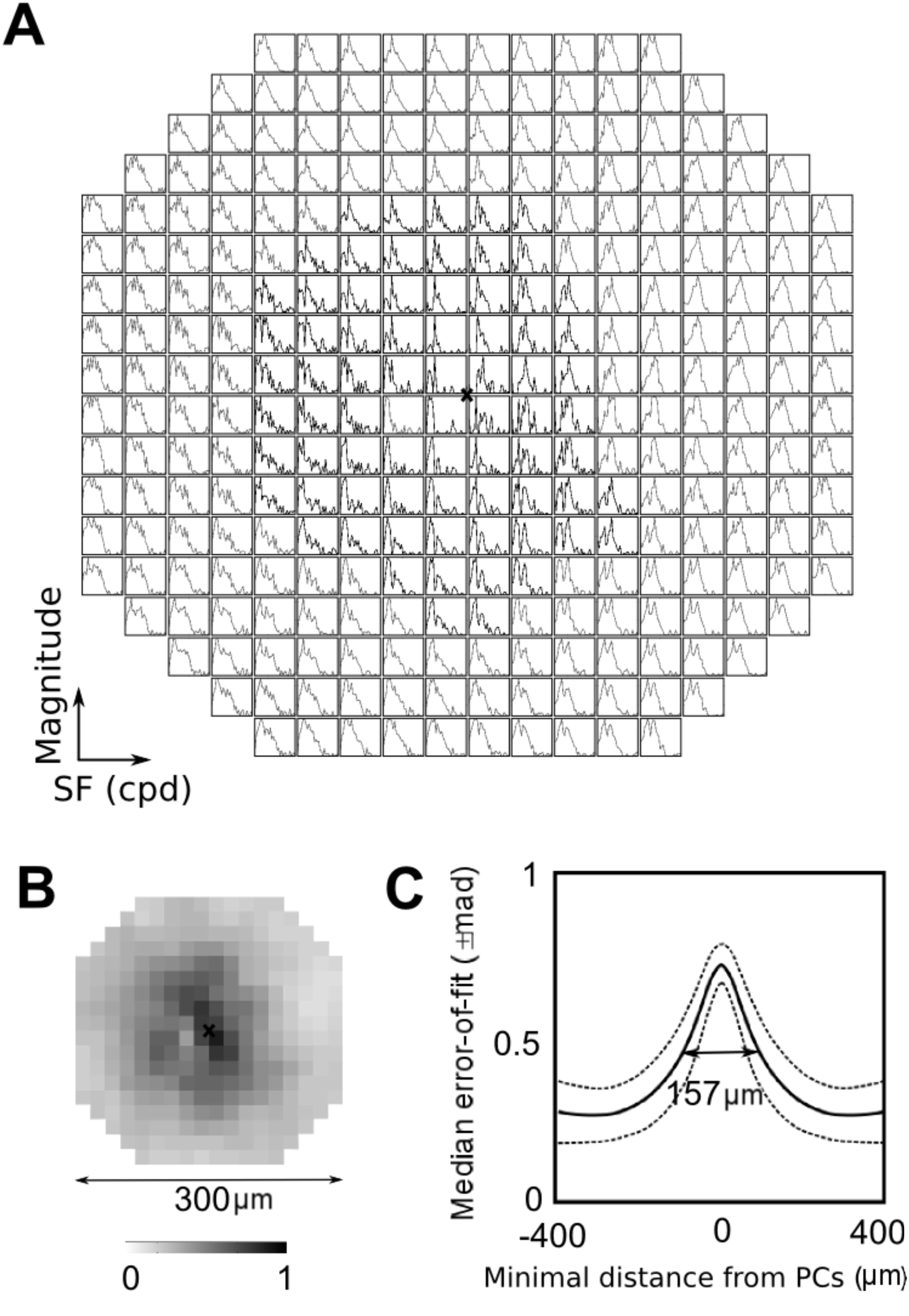
Spatial resolution of the SF maps. (A) Example of tuning curves in response to the 30 stimulus SFs around the PC circled in Fig. 1A2 and 1B2. Each tuning curve corresponds to one pixel of around 15 × 15 μm^2^. Tuning curves in black (gray) have an error-of-fit > 0.5 (< 0.5). Black cross is the PC location. (B) Error-of-fit around the same PC as in (A). (C) Median error-of-fit (± mad) with respect to the minimal distance from the PCs. Baseline level was around 0.27. The curve peaked at 0.72.

### 2.6 Existence of a local maximum and/or a minimum in the SF map

We then studied in more detail the SF representation around PCs. This requires to precisely locate the PCs, which was done automatically from the OR map, and to determine a typical scale for the PC’s vicinity in the SF map. Indeed, our previous results (Ribot et al., 2013) have indicated that cortical regions in the close vicinity from PCs have a high error-of-fit, typically exceeding 0.5 (See example in Fig. 2A, black tuning curves). This may be due to the spatial resolution of intrinsic optical imaging: In the literature, different values have been reported, ranging from 50μm (Grinvald et al., 1999) to 250-300μm (Orbach & Cohen, 1983; Polimeni et al., 2005). Defining a local scale around each PC thus requires that a sufficiently large number of pixels with well-defined preferred SF could be mapped. This issue is particularly important when mapping singularities or cortical locations with large gradient representation such as the dipoles reported here.

As shown in Fig. 2B, the error-of-fit seems to decrease with increasing distance from a PC. This was quantitatively confirmed by plotting the median error-of-fit following DOG interpolation as a function of the minimal distance from PCs (Fig. 2C). The full-width at half height for this curve was around 157 μm. In order to estimate the SF representation at cortical areas where the impact of the spatial resolution would be weak, we thus considered a circular region centered on each PC, and set the diameter of the region of interest at around twice this value, i.e. 300 μm. Pinwheel centers with overlapping regions of interest were rejected since this could cause a mutual interference in SF representation.

A pixel in the SF map was then defined as a maximum (or minimum) if its value belonged to the upper (or lower) SF values calculated over a hypercolumn. Different thresholds for defining the upper and lower bounds are presented in the manuscript (Fig. 6). The size of a hypercolumn was defined based on our previous results (Ribot et al., 2013): the SF structure in A17 is quasi-periodic, with periods ranging between 0.79 mm in the mediolateral direction and 1.43 mm in the anteroposterior direction. We estimated here a hypercolumn as a circle of 1.11 mm diameter (i.e. the average of the two values). In A18, the diameter of a hypercolumn was set to 1.5 mm (Ribot et al., 2013).

The results are illustrated in Figure 3 around two PCs (Fig. 3A). The corresponding hypercolumn is the region within the black circles in Fig. 3B, within which the distribution of preferred SFs appears in Fig. 3C. Based on this distribution, the two thresholds (dashed lines, denoted υ_LOW_ and υ_HIGH_) were calculated in order to define the upper and lower SF values (upper and lower quartiles in this example).

**Figure 3.**
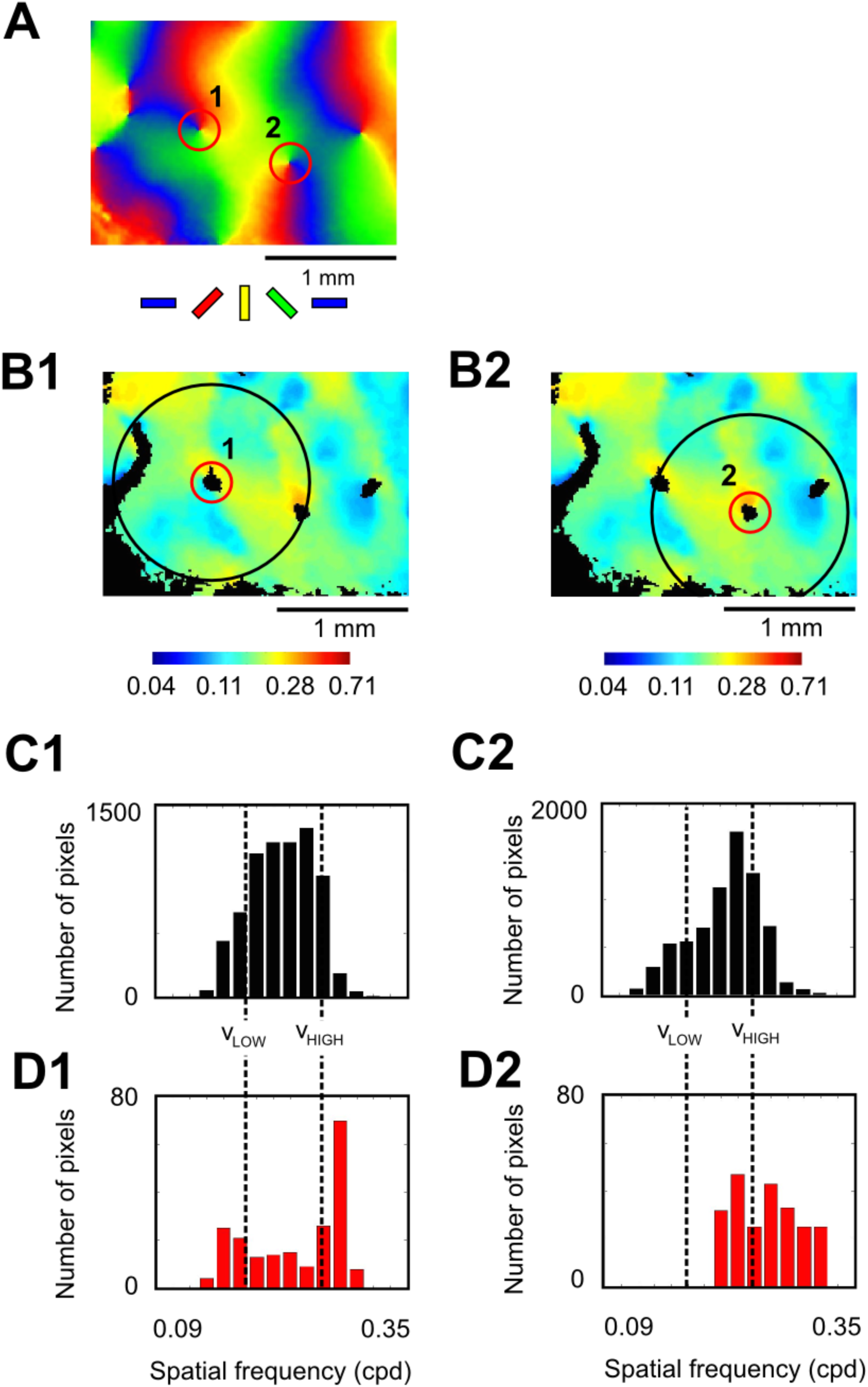
Methods for defining a semi-global extrema near a PC. (A) OR map in A18. Two PCs are selected (red circles). (B1) and (B2) SF map. The black circle corresponds to a SF hypercolumn centered on PC 1 and 2, respectively. (C1) and (C2) SF distribution within a hypercolumn centered on PCs 1 and 2, respectively (black circle in (B1) and (B2)). For each distribution, dotted lines correspond to the two thresholds νLOW and νHIGH defining respectively the lower fourth (25%) and upper fourth (25%) SF values. (D1) and (D2): SF distribution near PCs 1 and 2, respectively (red circle in (A), (B1) and (B2)). We observe that in the vicinity of PC 1 the SF representation takes values both in the upper and lower fifth quantiles shown in (B1). For this threshold, PC 1 thus has two semi-global extrema. On the other hand, PC 2 does not have any semi-global miminum.

Figure 3D shows the distribution of preferred SFs within 150 μm from the two PCs (Fig. 3A-C, red circles). The first PC had SF values beyond the two thresholds and thus had both a local maximum and a local minimum. On the other hand, the second PC only had SF values above the upper threshold, and thus had only a local maximum.

### 2.7 Geometrical description of a Pinwheel-Dipole

In order to ensure that the SF representation near PCs resembles a dipole, we developed an analytic model for fitting SF representation around PCs.

For this purpose, we considered a circular region of radius *R* = 150 μ*m* (the pinwheel area, PA) around a PC. The OR map can be classically described (Petitot, 2008) in the PA as the half angle of a complex coordinate *z* = *r*. exp (*i*. φ) (Fig. 4A, B), defined modulo π:

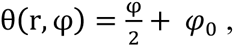

where *φ*_0_ is an arbitrary phase depending on the pinwheel considered.

**Figure 4.**
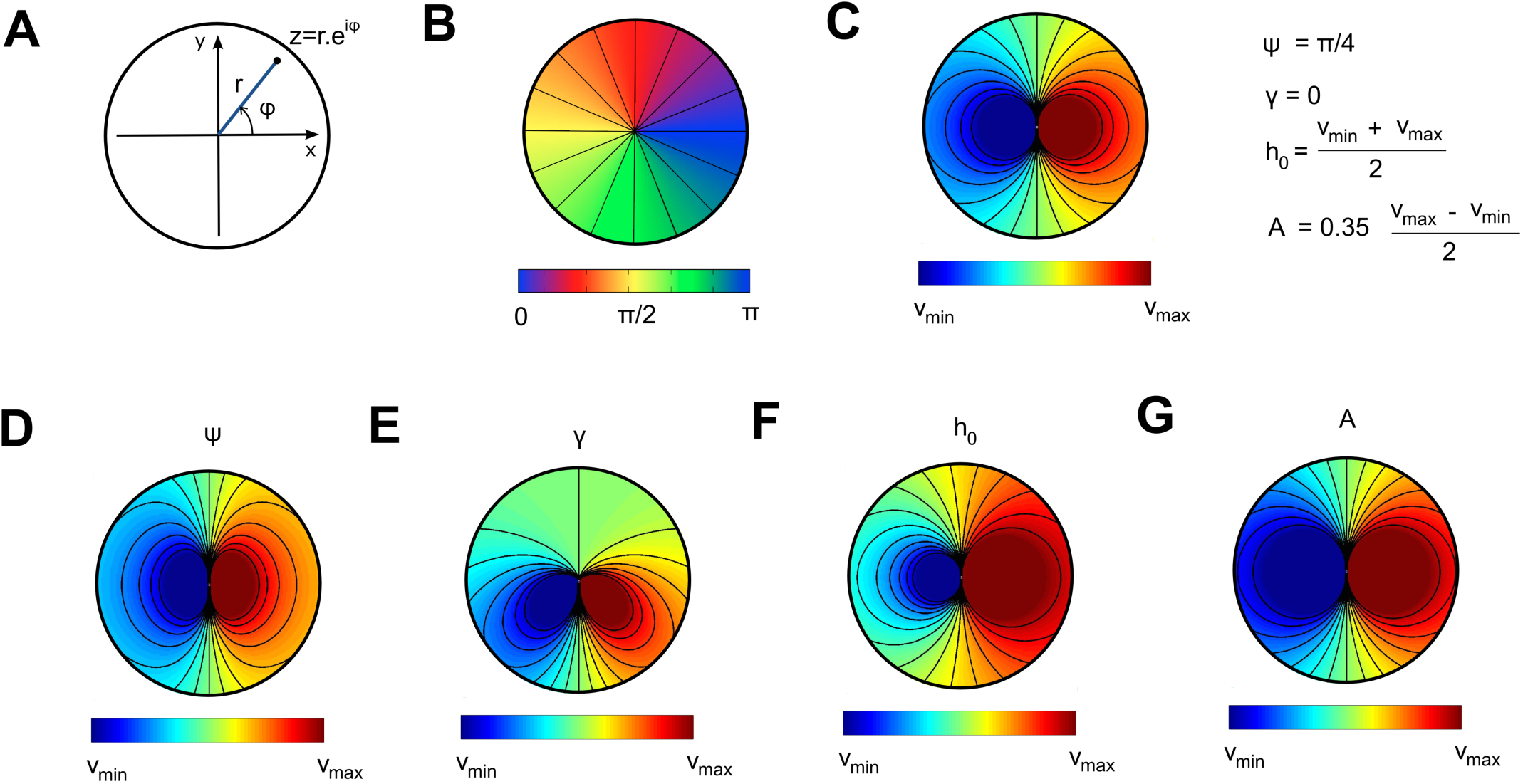
Geometric model of the pinwheel-dipole around PCs (A) Representation of a complex coordinate *z* = *r*. exp (*i*. φ). (B) Model of the OR PC (iso-OR lines in black). (C) Model of the SF dipole, with parameters equal to the average values across all fits (see Fig. 8). (D) to (F) Modifications of the dipolar architecture shown in (C) when varying the parameters ψ, γ, *h*_0_ and *A* in Equation (3).

As for our ideal model of dipolar topology, we used the classical two-dimensional electric potential created by an electric dipole, which is known to physicists to be proportional to the inverse of the distance from the center of the dipole:

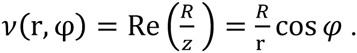

where *R* is a scale factor. This function exhibits a singularity at the PC (*z=0*) where it reaches both its minimal and maximal value and shows symmetrical circular level sets 180° apart from the singularity.

In order to fit this function with optical imaging data, we allowed deformations of this idealized model. The geometrical model of the SF map structure in the PA is defined as follows:

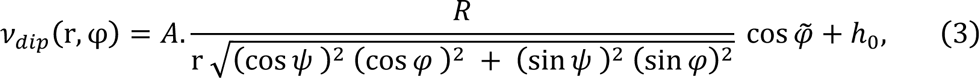

where 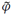 is an angular deformation allowing architectures in which circular level sets are not 180° apart:

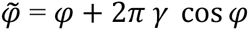

In addition, we considered equation (3) saturating at the maximum and minimum SF values *ν*_*max*_ and *ν*_*min*_. This saturation allows consistency with the experimental data by avoiding the divergence of *ν*_*dip*_ at the singularity.

Figure 4C-G display the architecture of these maps for different representative values of *ψ*, γ, *h*_0_ and *A*. The variation of these parameters allows continuous deformations of the shape of the iso-SF lines. It is important to notice, however, that the topology of the level sets remains unchanged.

### 2.7 Fitting the dipolar model

Equation (3) was fitted via a least-square method to the experimental data for those PCs having SF values in the upper and lower third of SF values calculated over a hypercolumn. Moreover, in order to have enough pixels for interpolating our dipolar model, we considered only PCs whose indeterminate zone (with an error-of-fit >0.5) covered less than 75% of the total area of interest. This corresponded to 115 PCs in A17 and 109 PCs in A18.

For each fit, the coefficient of determination R^2^ was calculated using the Matlab® function *regress* in order to have information on the goodness of fit of the model. Only those fit with R^2^ > 0.8 (i.e. fits for which 80% in the response variable can be explained) were kept in order to provide biologically relevant parameters in Fig. 8.

## 3 RESULTS

### 3.1 Spatial Frequency representation near Orientation Pinwheel Centers

A detail of OR and SF maps obtained with these experimental and image processing techniques is shown in Fig. 5 (see also Video S1 and Fig. S1). Four PCs are selected in A17 to illustrate the SF structure at these locations (Fig. 5A-B, black circles). The activity induced by the stimulation with various SFs within a 150 μm radius of PC 1 is shown in Fig. 5C. The immediate neighborhood of the PC is activated by a broad range of SFs. As the stimulus SF is increased, the activity, initially restricted to the lateral part of the PC, gradually invades the whole cortical area and almost completely fills it for SFs between 0.4 to 0.6 cpd. It then decays on the lateral part of the PC, while higher activity remains on the medial part above 0.9 cpd. There were no responses for SFs larger than 2 cpd. As a consequence, preferred SF representation shows spatial variations winding around the PC with local minimum (low SF) in the lateral part opposite to a local maximum (high SF) in the medial part (Fig. 5D1). The response tuning curves around these two semi-global extrema are represented in Fig. 5E, top panel.

**Figure 5.**
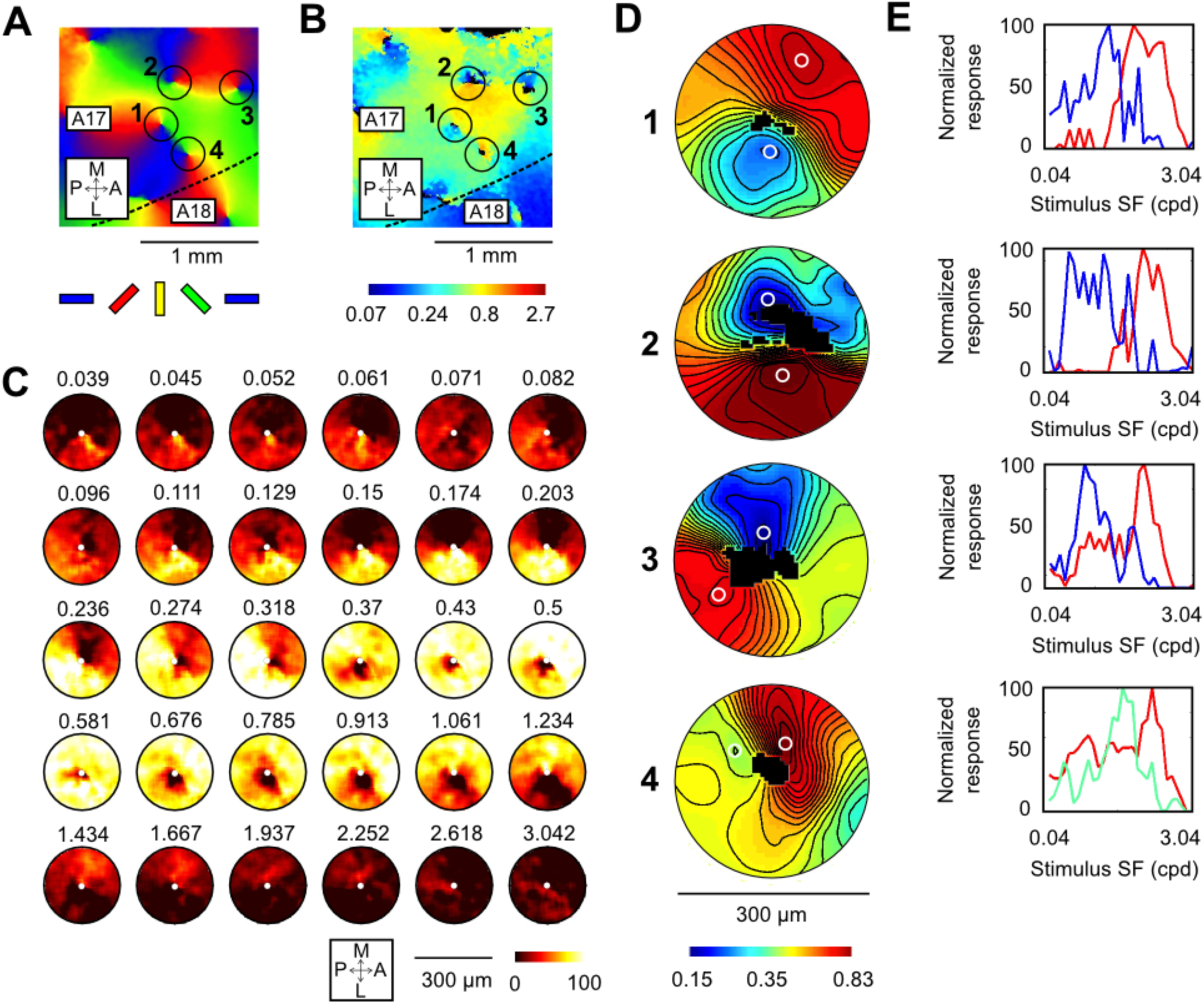
SF maps form dipoles at OR PCs. (A) OR map. Four PCs are selected (circles). Dashed line indicates the A17/A18 border. P: Posterior; A: Anterior; L: Lateral; M: Medial. (B) Corresponding SF map. Black pixels correspond to regions with error-of-fit > 0.5. Color map in logarithmic scale. (C) Maximal intensity maps around PC 1 for thirty SFs. Note the progression of the response from lateral to medial with increasing SF. (D) Zoom on the SF map for the four PCs circled in (A) and (B). An additional Gaussian low-pass filter with 35 μm sd was used to smooth the data and draw iso-SF lines (black). Logarithmic color map. For each panel, response tuning curve at two different locations (white circles) are represented in (E).

This structure constitutes a singularity in the SF map, co-localized with the PC. The SF structure for PCs 2 to 4 is shown in Fig. 5D2-4 and the response tuning curve for the two semi-global extrema are shown in Fig. 5E2-4. PCs 2 and 3 also exhibit maximum and minimum values separated by around 2.5 octaves. But while the extrema are aligned with PC 2, they are close to orthogonal in PC 3. The SF map of PC 4 has a clear maximum on one side of the PC, but no SF as low as for the three others on the other side of the PC. But it still displays a singularity, as indicated by the sharp transition in SF levels.

### 3.2 Existence of a local maximum and/or a minimum in the SF map around PCs

These examples have shown that extreme SF values seem to be represented near PCs. In order to validate quantitatively this observation, we identified semi-global extrema in the vicinity of each PC (circular region of interest with a 300 μm diameter, see Methods, Figures 2 and 3). We recall that semi-global maxima (or minima) correspond to values of the SF belonging to the last (or first) quantile of SF values calculated over a hypercolumn around the PC (see Methods, Fig. 3). They therefore depend on the quantile chosen.

The quantitative result of the analysis obtained on all cats is shown in Fig. 6. In this analysis, semi-global extrema were characterized as a function of the quantile chosen (Fig. 6A). Our results show that 89.6% (103/115) of the PCs in A17 and 76.2% (83/109) in A18 present the two semi-global extrema in the upper and the lower third of SF values found over a hypercolumn. Naturally, when the threshold is increased, the proportion of PCs with two semi-global extrema decreased. However, even for SF values in the upper and lower 10%, this percentage remained higher than the percentage for other configurations both in A17 (59/115=51.3%) and A18 (44/109=40.4%).

**Figure 6.**
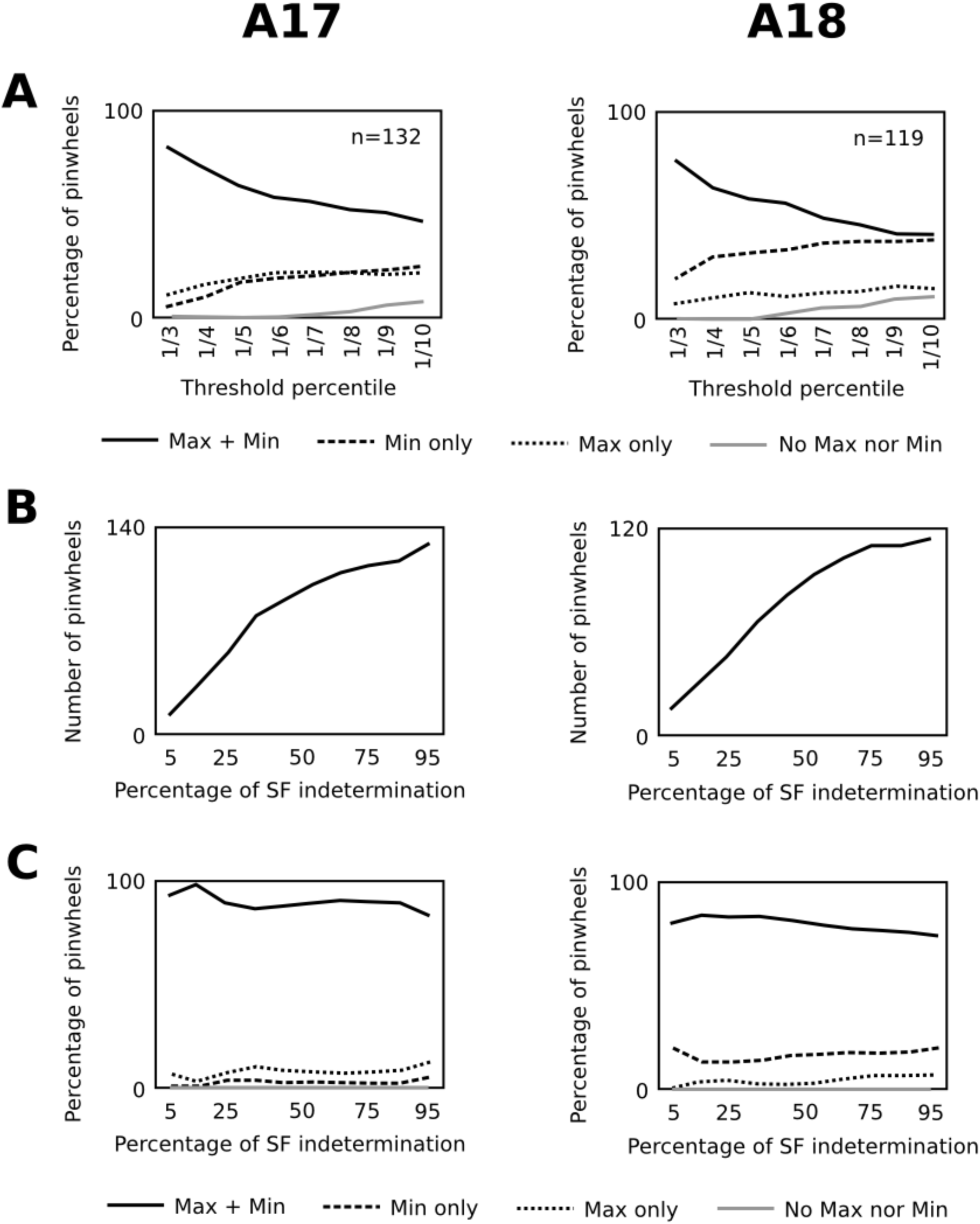
A majority of PCs has both a SF maximum and a SF minimum in their neighborhood. (A) Distribution of PCs in A17 (left) and A18 (right) with both a SF maximum and a SF minimum (black line), one single minimum (dashed line), one single maximum (dotted line) or not extremum (grey line) when changing the threshold percentile defined in Fig 3C. In this analysis, only PCs with a SF indeterminacy zone smaller than 75% were kept. (B) Cumulative distribution of PCs as a function of SF indetermination. (C) Distribution of semi-global extrema when changing the percentage of SF indetermination for PCs in A17 (left) and A18 (right). In (B) and (C), the threshold percentile was set to 1/3.

This threshold was then set at 1/3, and configurations were studied with respect to the percentage of SF indetermination (error-of-fit > 0.5) in the circular region of interest (Fig. 6B-C). Figure 6B represents the number of PCs in A17 and A18 with less than a given percentage of SF indetermination, and Fig. 6C the different configurations for these PCs. We found little impact of this parameter on the actual percentage of PCs with both a SF maximum and a SF minimum within 150 μm in A17 (> 80%) and in A18 (> 70 %).

We conclude that, in the vicinity of the PCs, the SF representation contains extreme values of the SF in a hypercolumn, and that a majority of locations exhibit the presence of both semi-global maximum and minimum. We further note that these results may challenge a model for which a single SF extremum would be located at the PC position (Shoham et al., 1997; Issa et al, 2000; Issa et al., 2008). In order to represent both semi-global maximum and minimum in a small neighborhood of the PC, the SF map may display a very specific organization. We now focus our attention on the fine organization of the SF map at PC.

### 3.3 Geometrical description of a Pinwheel-Dipole

Typical profiles of the SF map, as shown in Fig. 5, are familiar to physicists. The SF representation seems indeed analogous to the *dipolar architecture* displayed, for instance, by the electric potential created by an electric dipole. In order to ensure that the SF representation near PCs is indeed dipolar, we propose an analytic model for fitting SF representation around PCs (See Methods, Fig. 4).

Figure 7 shows an example of fit with this model in a region of interest covering A17 and A18. Only PCs with SF values in the upper third and lower third of SF values defined within a hypercolumn, and with a percentage of SF indetermination less than 75% are considered for these fits (Total: n=103 in A17 and n=83 in A18; White circles in Fig. 7A and 7B; Others appear in gray). PCs are numbered with decreasing coefficient of determination following the fit. A larger view of the SF representation around those PCs is shown in Fig. 7C, top rows. SF representation was rescaled so that red and blue colors corresponded to the local maximum and minimum of each PC, respectively. The fit with equation (3) is shown below by optimizing in each case the scalar parameters *ψ*, *γ*, *h*_0_, *A*, *ν*_*max*_ and *ν*_*min*_.

**Figure 7.**
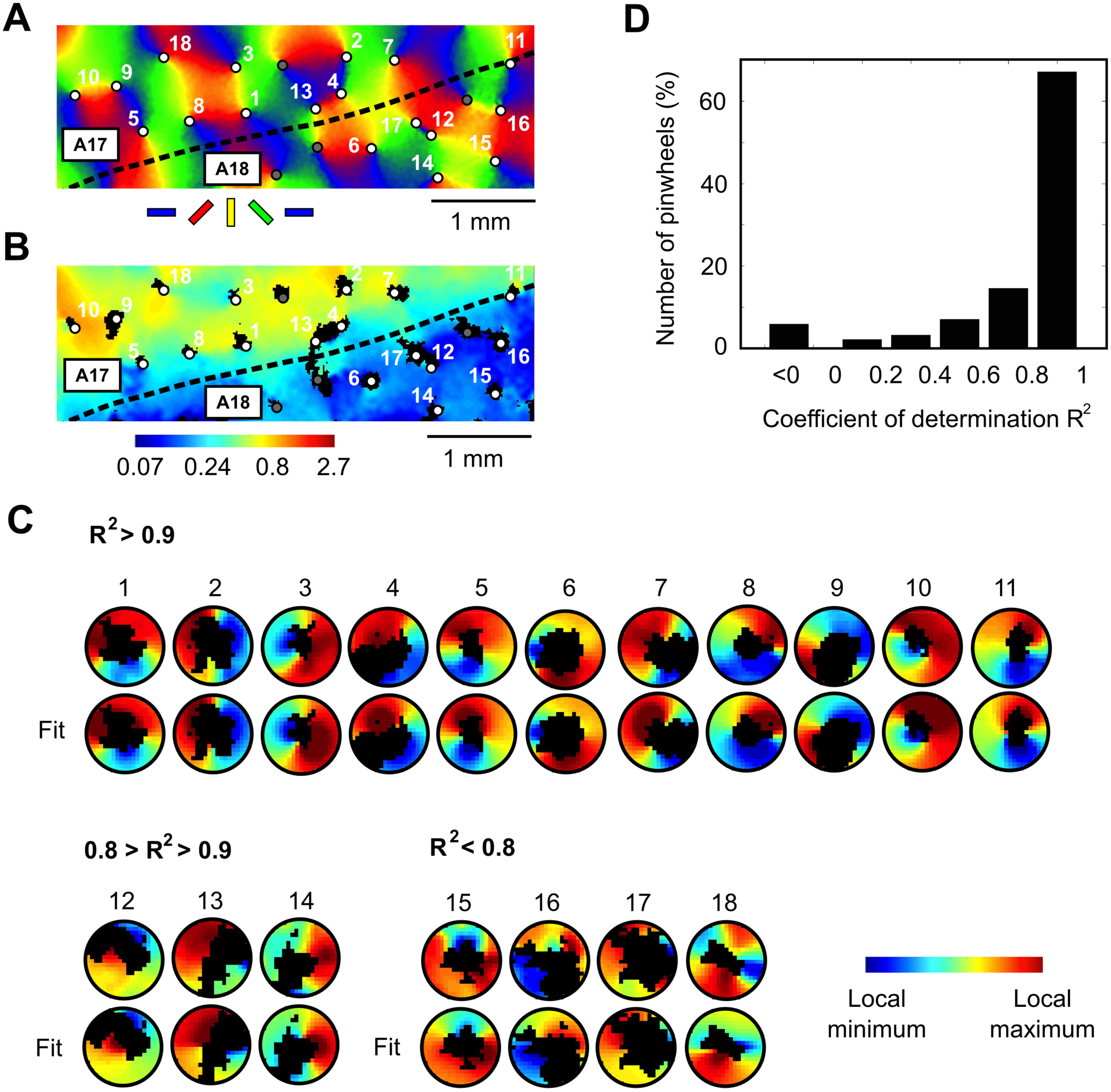
Fit of the SF representation around PCs with the dipolar model. (A-B) Maps of preferred OR and SF respectively. Scale bar: 1 mm. OR pinwheels indicated by the white and gray filled black circles. Gray PCs (not fitted) correspond to those whose SF map either have only one extremum (Threshold percentile = 1/3) or the indeterminacy region covers more than 75% of the 150 μm circle. (C) Fits of the SF representation of the 18 PCs (white dots in (A) and (B)) with the dipolar model in Equation (3). The least-square errors fits are shown in the lower rows and experimental data in the upper row. They are displayed with decreasing coefficient of determination (R^2^) order following linear regression between modeled and experimental data. Note that R^2^ can be negative, as the linear regression performed in Matlab® (function *regress*) was done without any DC component (the DC component h0 in equation (3) was optimized beforehand). (D) Distribution of R^2^ among the 186 PCs tested.

This test confirmed that a majority of PCs in the two visual areas show a dipolar SF organization. In detail, among the 186 PCs tested, we found that around two third of the fits had a coefficient of determination R^2^ greater than 0.8 (Fig. 7D, 123/186=66.1 %), i.e. that 80% of the variations could be captured by the model. Fits with R^2^ less than 0.8 typically corresponded to complex SF representation with multiple minima and/or extrema. No difference could be found between A17 (68/103=66.0%) and A18 (55/83=66.3%).

### 3.5 Statistics of the fits

Figure 8 shows the distribution of the parameters in equation (3) that were obtained from the fits over the 123 PCs with R^2^ greater than 0.8. This led to a robust estimate of each parameter in the model.

**Figure 8.**
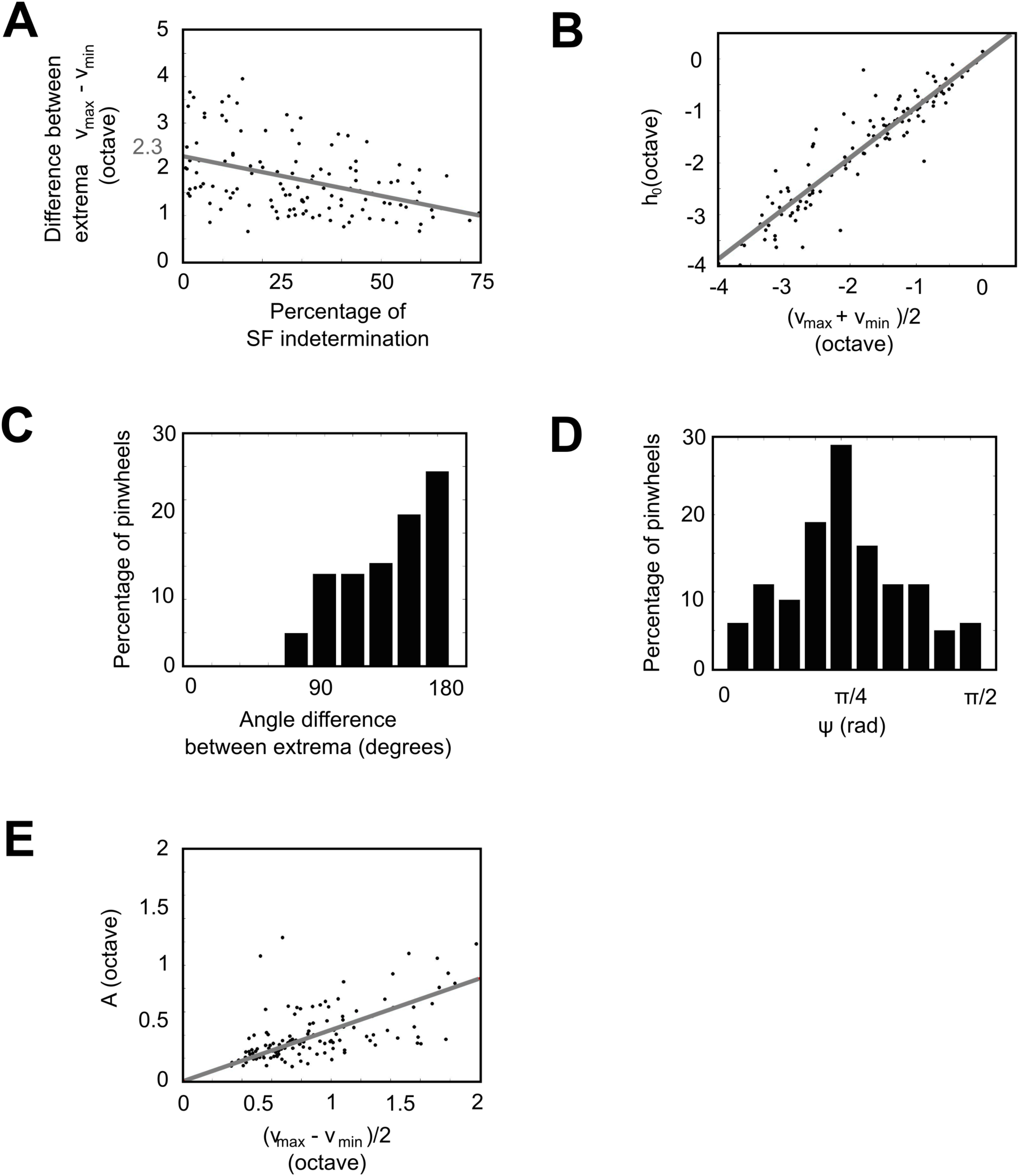
Statistics of the fits. Only those fits with R^2^ > 0.8 were kept. (A) Estimate of the difference between νmax and νmin. This difference is plotted with respect to the percentage of the surface of SF indeterminacy within 150 μm. Linear regression is shown in grey. (B) Scatter plot of h0 with respect to (νmax+ νmin)/2. Linear regression is shown in grey. (C) Distribution of the angle between the maximum and the minimum (estimated from the γ parameter) with respect to the PC. The peak at 180 degrees shows that extrema tends to be aligned with the PC. (D). Distribution of ψ. (E) Scatter plot of *A* with respect to (νmax-νmin)/2. Linear regression is shown in grey.

First we evaluated how the difference between the local SF extrema *v*_*max*_ − *v*_*min*_ was distributed among the PCs. For this purpose, *v*_*max*_ − *v*_*min*_ was plotted with respect to the percentage of SF indetermination (Fig. 8A). We found that the difference *v*_*max*_ − *v*_*min*_ decreased with the percentage of SF indetermination (slope of -1.67) and crossed the ordinate axis for 2.28 octaves. This suggests that extreme SF values tend to be hidden when the zone of indeterminacy is large. This is consistent with the idealized dipole model that locates extrema at the singularity: the smaller the indeterminacy area, the larger the SF range. Eventually, we conclude that the mean (± sd) value for *v*_*max*_ − *v*_*min*_ in equation (3) is 2.28 ± 0.81 octaves.

Then, the DC component h_0_ was plotted with respect to the average value between *v*_*max*_ and *v*_*min*_ (Fig. 8B). We found a noticeable correlation between the two parameters (slope 0.99). This indicates that the two semi-global extrema tend to be of equal size (see Fig. 4F). An estimate (± sd) of the coefficient h_0_ in equation (3) is thus 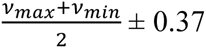 octaves.

Figure 8C shows the angle difference between the two extrema, which is related to the parameter γ in equation (3) (see Figure 4E for example). Remarkably, we found that this angle was biased toward the highest values, indicating that the location of the maximum and the location of the minimum are preferentially located symmetrically with respect to the PC. Variations however existed, and the sd of the angle was evaluated to 56 degrees.

Finally, we evaluated the values of the parameters ψ and A. The mean (± sd) value for the slope ψ was 0.7 (± 0.37) radians (Fig. 8D), i.e. close to π/4 (∼ 0.785) for which no deformation of the topology occurs compared to the idealized dipole model. The parameter A was estimated by plotting its values with respect to (*v*_*max*_ − *v*_*min*_)/2 (Fig. 8E). We found a positive correlation between these two parameters (slope = 0.44). An estimate (± sd) of the parameter A is thus 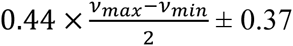 0.37 octave.

The resulting dipole for the average values of the parameters *h*_0_, γ, ψ and A is the dipole represented in Fig. 4C.

## 4 DISCUSSION

These results thus shed new light on the structure of cortical maps in cat early visual cortex. We show that singularities in the OR map form dipoles in the SF map for at least 54.9% (123/224) of the PCs tested in A17 and A18. Other PCs couldn’t be accurately fitted with the dipolar model we proposed (63/224=28.1%) or did not have simultaneously a maximum and a minimum in the upper and lower third of SF values defined within a hypercolumn (38/224=17%). Even in the latter case, however, SF discontinuity can still occur at pinwheel singularities (See Fig. 5D4 for instance), with a single extremum located close to the PC but not at the center of it.

### 4.1 At which spatial scale pinwheel-dipoles appear?

In order to exhibit the structure of the SF map near OR PCs, one needs to use a neuroimaging technique at a scale sufficiently macroscopic to identify SF preference over a large number of PCs, and fine enough to resolve spatial details of the topology. Optical imaging of intrinsic signals satisfies these requirements and was used in this study.

First, our aim was to map a sufficiently large number of PCs in order to statistically ensure the reproducibility of the SF structure at these locations. Intrinsic optical imaging enables mapping large portions of the early visual cortex, containing several dozens of pinwheel locations. In contrast, more recent techniques with single cell resolution (Ohki et al., 2006) only allow mapping a surface on the order of two to three PCs simultaneously (Nauhaus et al., 2013). Second, previous studies based on electrophysiology have concluded that preferred SF is only weakly clustered among neighboring neurons in the visual cortex of felines (DeAngelis et al., 1999) and bush babies (Purushothaman 2009). Our own electrophysiological data (see Fig. S2), although consistent with a sharp transition in the SF map at crossing OR PCs, supports this weak clustering. Imaging SF maps in cat with single cell resolution is thus not likely to be suitable to evidence a collective organization. Intrinsic optical imaging combines the two advantages of recording the activity induced by a population of neurons for a wide cortical surface, thus revealing a clear SF map over a large number of PCs.

Notwithstanding, optical imaging has a relatively modest spatial resolution. According to previous reports, the spatial resolution of instrinsic optical imaging lies between 50μm (Grinvald et al., 1999) and 250-300μm (Orbach & Cohen, 1983; Polimeni et al., 2005). As a consequence, the signal recorded at each cortical location could be influenced by signals from neighboring neurons. Near SF dipoles where extreme values of the SF are represented, this leads to the superposition of signals from pixels having a wide range of preferred SFs. Accordingly, we found that the error-of-fit following DOG interpolation was maximum at PC location. However, at around 80 μm from PCs, the error-of-fit was already half weaker, and setting the radius of the region of interest at 150 μm was sufficient to map different ranges of preferred SFs (See example in Fig. 2) and to reveal a clear but non-trivial topology.

### 4.2 Dipoles and uniform coverage

The SF dipoles create discontinuities in SF representation as the OR PCs create sharp gradients in the OR map. This result is reminiscent of early investigations by Das and Gilbert (1997) on the relationships between the map of visual space and the OR map. The authors showed that the map of visual space had local distortions that also registered with OR PCs. Although these results were later disputed (Hetherington & Swindale, 1999; Buzás et al., 2003), they raised fundamental questions about how multiple cortical maps should be represented on a single cortical sheet.

At the present time, the dominant theory proposes that continuity and coverage uniformity constitute fundamental principles governing the organization of these maps (Swindale 1991; Swindale et al., 2000). This view is supported by several experiments showing that lines of iso-properties tend to run perpendicular to each other (Yu et al., 2005; Nauhaus et al., 2012), and that OR PCs are situated preferentially at local SF extrema (Shoham et al., 1997; Issa et al, 2000; Issa et al., 2008).

The dipolar structure reported here clearly does not fit with this theory, since a specific range of ORs is represented for a specific range of SFs. On the other hand, the dipole ensures the representation of a large range of SFs in the immediate neighborhood of the singularity, as the pinwheel does in the OR map. As shown in a companion paper (Romagnoni et al., 2014), this could be highly beneficial for visual perception. Indeed, with a single SF represented at a PC, the pinwheel would be blind for a large range of SFs. On the other hand, with a pinwheel-dipole singularity, a much larger range of the visual attributes would be processed simultaneously, yielding a balanced estimate of the OR and the SF contained in a stimulus.

### 4.3 Dependence between orientation and spatial frequency representations

The striking co-localization of OR PCs and SF dipoles likely emerges from developmental constraints. Indeed, immature OR maps exist early after eye opening (Crair et al., 1998) at a time when only low SFs are represented (Tani et al., 2012). Afterwards, both maps jointly develop with visual experience (Tani et al., 2012). Such interdependence of the two features is also reflected by the variations of preferred SF with stimulus OR (Webster & De Valois, 1985; Issa et al, 2000), and by the response to complex stimuli when varying SF, speed and direction (Basole et al., 2003). We conclude that OR and SF are not independently represented in cat primary visual cortex, and therefore should be studied jointly.

Whether this dependence between maps is also true for other mammals, including humans, however remains undetermined. It would not be surprising if different species adopted distinct coding strategies adapted to the number of features encoded in the primary visual cortex and to their specific circuit organization. For instance, maps of preferred OR and SF in the primate tend to show sharp orthogonality (Nauhaus et al., 2012), which is not the case of ferrets (Yu et al., 2005). However, these global statistics on the whole maps shall be finely studied near PCs to uncover SF representation at these locations.

### 4.4 Exhaustivity and parsimony at pinwheel-dipole centers

Pinwheel and dipole structures have fascinating and universal geometric features. As shown in the companion paper (Romagnoni et al., 2014), the pinwheel is the unique continuous architecture ensuring exhaustive representation of a periodic quantity with minimal redundancy. Similarly, the dipole is the most parsimonious continuous topology ensuring a local representation of a non-periodic range of SF values. Both topologies thus allow collecting economically all information, which could be fundamental for visual processing. In particular, neurons at PCs have been shown to exhibit properties distinct from those in the regular domains: they show strong resistance to monocular deprivation (Crair et al., 1997), are maximally sensitive to OR adaptation (Dragoi et al., 2001) and have a high dynamical variability (Schummers et al., 2002). They also receive intracortical inputs from domains signaling a wider range of ORs (Yousef et al., 2001, Schummers et al. 2002). Properties related to SF perception at pinwheel-dipole centers, such as enhanced SF adaptation (Sharpee et al., 2006) or their interactions with nearby cortical regions, could exhibit similar specificities, and, for these purposes, collecting a large range of SFs would be beneficial.

## 5 CONCLUSION

In summary, we show that a majority OR PCs forms SF dipoles in cat early visual cortex. The dipolar organization is not trivial and strongly differs from pioneering observations showing a single SF extremum located at PC locations. Pinwheel-dipole co-localization also argues against uniform coverage hypothesis, according to which all combinations of the attributes should be uniformly represented in the cortex. Instead, each single topology represents locally a large range of visual attributes. This specificity may provide indications on the general principles at play in the mutual arrangements of cortical functional maps as well as their possible role in visual perception.

## ACKNOWLEDGMENTS

We thank Timothé Coulais for animal care and Sidney Wiener for discussions and comments on the manuscript.

## ABREVIATIONS

OR: Orientation
SF: Spatial frequency
PC: Pinwheel center
A17: Area 17
A18: Area 18

## AUTHOR CONTRIBUTIONS

J.R. designed and performed experiments, analyzed experimental data and wrote the paper. A.R. analyzed experimental data, designed the model and wrote the paper. C.M. provided administrative and financial support for the experiments, designed and built the experimental post. D.B. and J.T. designed the model, supervised the project and wrote the paper.

## SUPPLEMENTARY MATERIAL

**Video S1.** Single-condition maps for each SF of stimulation are shown sequentially for the same cat as in Fig. 5. Each panel corresponds to one SF of stimulation, as in Fig. S1. Each frame corresponds to one stimulus OR rotating at a frequency of 2 rpm and drifting at 2Hz. Circles correspond to PCs defined in the OR map. The speed of the movie is twice the speed of rotation.

**Figure S1.**
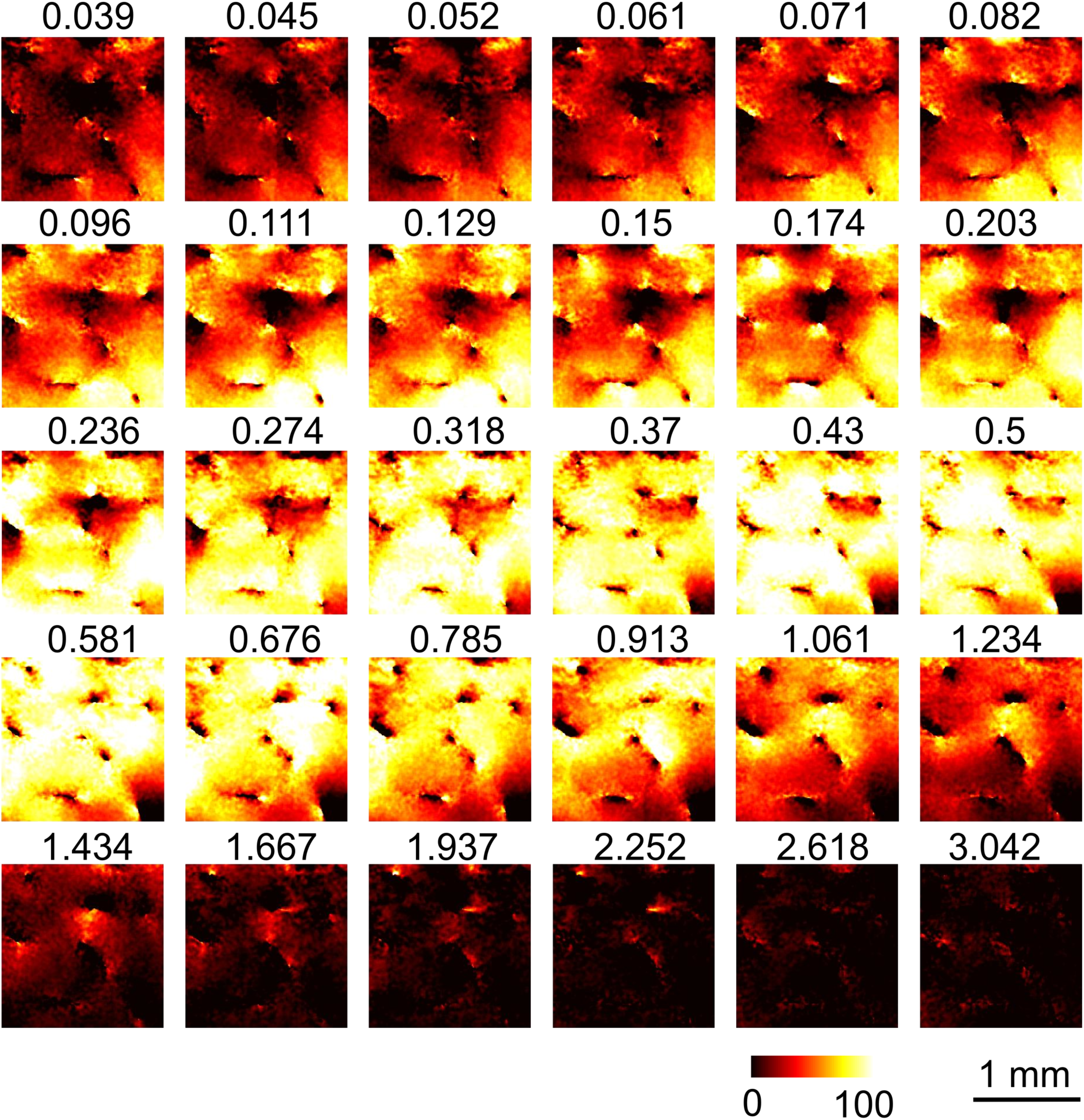
Maximal intensity maps for the animal shown in Fig. 5 and Video S1. Thirty SFs were presented to the animal. The signal at each pixel that elicited the maximum activation was normalized at 100%. Scale bar: 1 mm.

**Figure S2.**
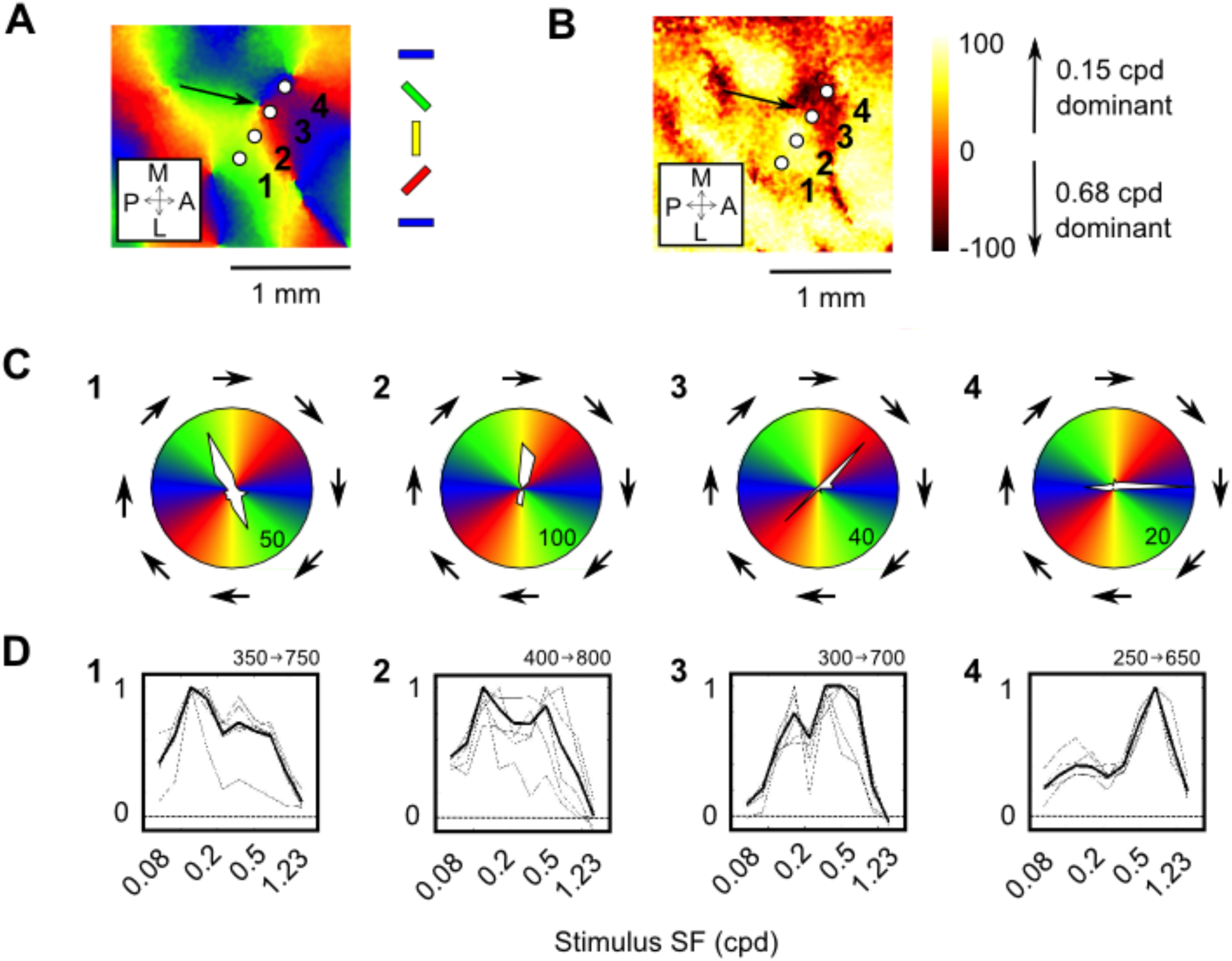
Electrophysiological investigation of SF representation near PCs. Detailed method for electrophysiology are described in Text S1. (A) OR map with four electrode penetrations (white dots) performed around a PC (black arrow). (B) Difference between maximal intensity maps induced for stimulus SF of 0.15 cpd and 0.68 cpd. Domains in yellow-white elicit a larger response for 0.15 cpd, those in red-black for 0.68 cpd. The PC falls at the border between two patches. (C) Polar plots of firing rate (Hz) induced by the presentation of eight ORs for these four penetrations. Arrows indicate the direction of movement of the stimuli. The maximum spike rate is indicated in the bottom right corner. (D) Corresponding SF response curves for stimuli at the preferred OR. X-axis scale is logarithmic. Thin gray lines are normalized firing rate at five depths in superficial layers separated by 100 μm. Minimum and maximum depths are shown at upper right. Thick black line is total normalized spike rate across depths. The zero baseline corresponds to spontaneous activity.

While preferred orientation estimated most superficially (Fig. S2C) matched the one calculated with optical imaging, the SF tuning exhibited fluctuations among the different depths (Fig. S2D, thin lines). This possibly reflected the weak clustering of neurons for SF found in previous studies (DeAngelis et al., 1999, Purushothaman 2009). Although it was generally broadly tuned, the averaged activity (thick line) confirmed the variation in preferred SF observed with optical imaging (Fig. S2B). Locations 1 and 2 prefer SFs lower than 0.2 cpd. Location 3 (within 100 μm of the PC) yields maximal response between 0.37 and 0.68 cpd (Fig. S2D3). On the other side of the PC (location 4), peak response is for 0.68 cpd.

Methods. Optical imaging: two SFs of 0.15 cpd and 0.68 cpd were used to reduce the duration of the recording (10 minutes) and data were processed *off-line*. Targeted multi-unit recordings were made using the NeuraLynx system after off-line analysis of optical imaging data. Tungsten microelectrodes (1–2 MΩ at 1 kHz) were used. The electrical signal was amplified and band-pass filtered between 0.3 and 3 kHz. Then, unit responses were thresholded with a window discriminator to separate spike activity from electrical noise. The average depth from cortical surface for the first recording was 325 ± 65 μm s.d. across the four penetrations. Sine-wave gratings of different ORs (n=8) were presented randomly to the animal. The gratings, with a SF of 0.37 cpd, drifted perpendicularly to the OR at a temporal frequency of 2 Hz. Sixteen directions were presented 20 times to the animal for a period of 1 s each, interleaved by blank-screen stimuli for 1 s in order to provide measures of spontaneous activity. Once OR preference was determined, subsequent stimuli consisted of pseudo-randomly ordered sine-wave gratings spanning a logarithmic range of SFs between 0.08 and 1.23 cpd (10 SFs in total, with 0.44 octave increments). The gratings were presented at the preferred OR, and 30 trials were presented for each stimulus. Blank-screen stimuli were presented for 1 s interleaved with grating stimuli to measure spontaneous activity. This spontaneous activity was subtracted from the activity evoked by each SF. Five recordings were made for each penetration at regular intervals of 100 μm. The average depth of the deepest recording was 725 ± 65 μm s.d. among the four penetrations. Then, spike activity was summed up among the five depths to obtain an estimate of the SF response of the population of neurons at each targeted location.

